# Grid Cell Percolation

**DOI:** 10.1101/2022.08.26.505489

**Authors:** Yuri Dabaghian

## Abstract

Grid cells play a principal role in enabling mammalian cognitive representations of ambient environments. The key property of these cells—the regular arrangement of their firing fields—is commonly viewed as means for establishing spatial scales or encoding specific locations. However, using grid cells’ spiking outputs for deducing spatial orderliness proves to be a strenuous task, due to fairly irregular activation patterns triggered by the animal’s sporadic visits to the grid fields. The following discussion addresses statistical mechanisms enabling emergent regularity of grid cell firing activity, from the perspective of percolation theory. In particular, it is shown that the range of neurophysiological parameters required for spiking percolation phenomena matches experimental data, which points at biological viability of the percolation approach and casts a new light on the role of grid cells in organizing the hippocampal map.

## I. INTRODUCTION AND MOTIVATION

Cognitive representation of space is sustained by the spiking activity of “spatially tuned” neurons, such as hippocampal place cells, head direction cells, parietal cells, border cells, and others [1, 2]. A particularly curious pattern of activity is exhibited by the grid cells in the rats’ Medial Entorhinal Cortex (MEC) that fire in compact domains centered at the vertexes of a triangular lattice, tiling the navigated environment [3] (Fig. 1A). The exact principles by which these cells contribute to spatial awareness remain a matter of debate. It is commonly assumed that MEC outputs are used to represent the animal’s ongoing location and to establish global spatial metrics [4, 5]. However, extracting these structures from the spike train patterns is a complex task: since the animal can visit one firing field at a time, the sequences of grid cell responses depend on the shape of the rat’s trajectory and can be highly intermittent. In absence of simple universal decoding algorithms, the effect produced by the grid cells in the downstream networks may depend primarily on activation frequency: persistently firing cells contribute most, while the ones that activate sporadically produce smaller impacts [6]. The maximal frequency of a given grid cell’s responses is achieved over periods when the animal runs through its firing fields in sequence, without omissions (Fig. 1A). Question is, under which conditions such contiguous firing can be produced in a given environment, how common are the regularly firing cells and so forth.

**FIG. 1.**
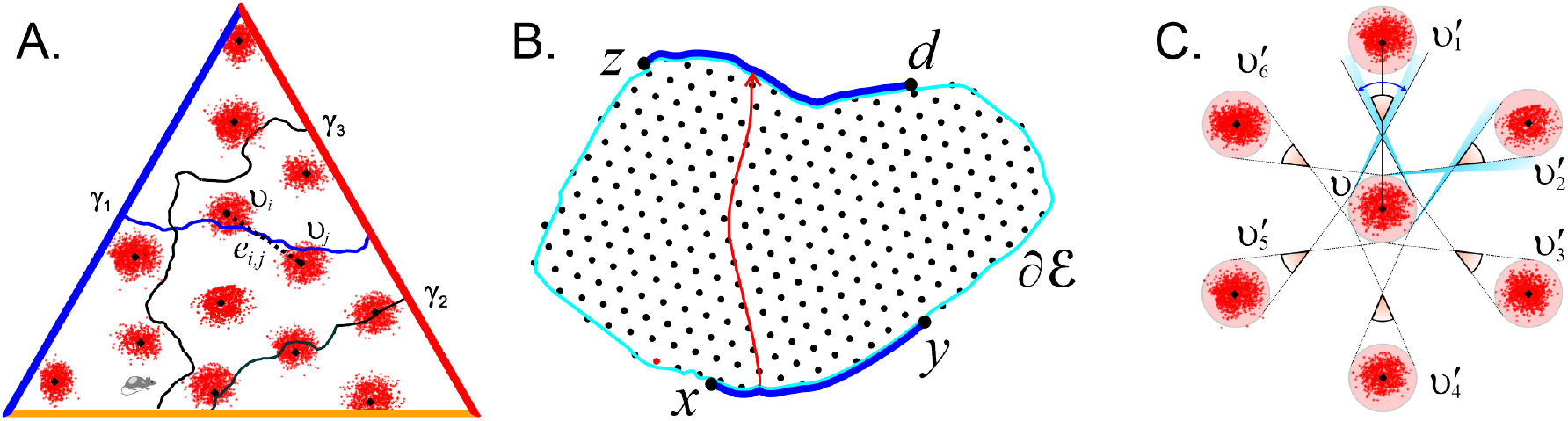
Grid cells. (**A**). Grid fields form a hexagonal lattice embedded into a 3×3 m triangular enclosure. Vertexes of the lattice are shown by black dots. The diameter of each field is comparable to the animal’s body size. Three curves, *γ*_1_, *γ*_2_ and *γ*_3_, represent paths segments extending from one side of a triangular environment to another. The path *γ*_1_ crosses the grid fields *υ*_*i*_ and *υ*_*j*_ eliciting spikes in both, thus opening the vertexes *v*_*i*_ and *v*_*j*_ and instantiating the edge *e*_*ij*_ between them (dashed line). Path *γ*_2_ percolates through the grid fields in sequence, without omissions, *γ*_3_ avoids grid fields. **B**. A percolation theory setup: a path extending from one side of a lattice domain *V*_*ℰ*_, a boundary segment ([*a, b*] ∈ ∂*ℰ*) to another ([*z, d*] ∈ ∂*ℰ*) may open vertexes or edges with fixed probabilities *p*_*v*_ and *p*_*e*_. **C**. The range of directions that lead from a grid field υ to its six immediate neighbors, *υ*_1_, …, *υ*_6_, are marked by pink. Directions along which a straight line escapes between the fields are marked by blue. For *ξ*_*g*_ = 1/2, the escape directions disappear.

Curiously, these questions are reminiscent of the problems addressed in percolation theory, which describes propagation of diffusive substances (liquids or gasses) through porous media. The key question addressed by the theory is whether a permeable domain *ℰ* allows diffusive leaks from one side of its boundary to another [7], i.e., whether the trickling through the pores can form uninterrupted sequences connecting the opposite sides, or *percolate*^1^ through, *ℰ*.

Of particular interest for the following discussion are mathematical models of percolation, in which physical media is represented by a segment of a regular lattice *V* enclosed within a domain *ℰ*. Depending on the setup, either the vertexes *v* or the edges *e* of the lattice *V*_*ℰ*_ represent the leaking pores, which may “open” or “close” with some fixed probabilities [7, 8] (Fig. 1B). A key result of the theory is that exceeding critical thresholds (for triangular lattices, 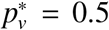 for the vertexes and 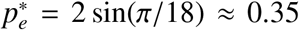 for the edges), marks the onset of a *percolating phase*, in which uninterrupted sequences of open sites, connecting opposite sides of the lattice *V*_*ℰ*_, become statistically common [9].

The analogy with the grid cells can be formalized as follows. Consider a triangular lattice 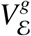 with vertexes centered at the firing fields of a cell *g*. A vertex 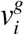 opens if the cell *g* fires at the corresponding field 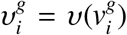. If the rat runs consecutively through two neighboring fields, e.g, from 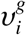 to 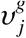 on Fig. 1A, eliciting spikes in both, then the edge 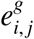 between them also opens. If a path *γ* induces a sequence of conjoint open edges,

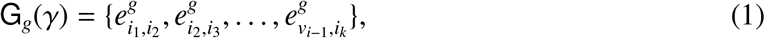

(and hence runs through a series of open sites 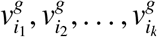), it will be said to percolate *g*. The spiking pattern of a cell *g* triggered by the rat’s moves can then be described by a sequence of open vertexes and edges, i.e., represented by discretized the path (1).

In the following, it is shown that, for a certain scope of firing parameters, the paths *γ* extending through *ℰ* may systematically percolate groups of grid cells, which may then play a particular role in representing spatial information. The activity of MEC network can therefore be studied from the perspective of identifying such high-impact cells, understanding their role in representing the navigated paths, testing whether the parameters required for percolation are physiologically viable and so forth.

## II. RESULTS

### Preliminary estimates

Grid cell percolation depends on the probability with which a generic trajectory runs into the grid fields and the probability of eliciting spiking responses. The former is controlled by the ratio between the field size *D*_*g*_ and the grid spacing *a*_*g*_,

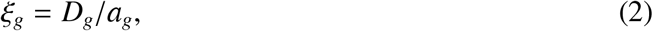

while the latter depends on the maximal firing rate *A*_*g*_ and the animal’s speed *s*.

*Lattice parameter ξ*_*g*_ defines the range of directions that lead from a grid field to one of its immediate neighbors. If the gap between fields is wider than the field size, *ξ*_*g*_ < 1/2, then a finite fraction of straight directions originating at a given grid field form “escape corridors”— passageways in-between the surrounding fields (Fig. 1C). At *ξ*_*g*_ = 1/2 such directions disappear, suggesting that *ξ*_*g*_ ≳ 1/2 is required for enforcing the percolation. However, this requirement must be strengthened further, for two reasons. First, the trajectory cannot not just brush on the field’s side, where the firing rate is too small—it should pass sufficiently close to the center, to induce reliable spiking responses. Second, the “empirical” size of a field is defined by the lengths of the typical paths that run though it, rather than the field’s diameter. A simple correction to (2) can hence be obtained by replacing the diameter *D*_*g*_ with the length of an average chord cutting through the field, 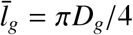 [10, 11], which yields *ξ*_*g*_ ≳ 2/π (Fig. 1B).

If *ξ*_*g*_ grows further, the angular domains leading from field to field begin to overlap. To maintain unambiguity of representation, the lattice parameter (2) should remain close the marginal value,

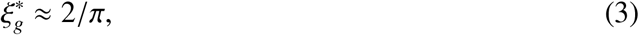

which matches the experimentally observed “isometric relation” [3, 13].

*The opening probabilities* on a grid field lattice depend on the parameters of neuronal activity and the animal’s moves. If the maximal spiking rate of a cell *g* is *A*_*g*_, then the mean rate is

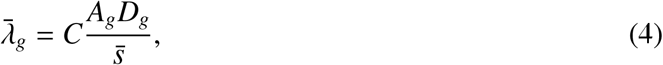

where *C* ≈ 0.06 is a geometric coefficient, and 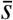 is the mean traversal speed (see Appendix, Sec. IV). Experimentally, the grid field sizes sampled along the ventro-dorsal axis of MEC range, in smaller environments, from about 10 cm to about 20 cm [3, 12], while the mean rates co-vary between *A*_*g*_ ≈ 21 Hz to *A*_*g*_ ≈ 11 Hz, i.e., the product *A*_*g*_*D*_*g*_ tends to increase as the field sizes grow. In larger environments, the firing rates remain approximately same, while field sizes cover the range 50 ≲ *D*_*g*_ ≲ 120 cm, driving 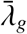 to higher values [14]. Thus, if the rat spends 2 − 3 seconds within a field, the expected firing probability,

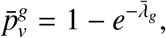

is high—the corresponding cell fires almost certainly. In particular, the mean vertex opening probability exceeds the critical value 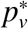, which suggests that vertex percolation may indeed take place in the parahippocampal network. On the other hand, the expected probability of opening an edge *e* can be estimated from

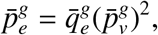

where 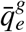 is the mean probability of reaching a grid field starting from its closest neighbor. Geometrically, 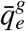 depends primarily on the lattice parameter *ξ*_*g*_ (Sec. IV). For the experimentally observed *ξ*_*g*_ ≈ 2/3, the estimated value is about 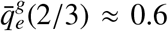, which, for high enough 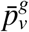, reaffirms the possibility that some grid cells may be systematically percolated during navigation.

### Grid field percolation

These hypotheses can be tested by simulating rat’s navigation in triangular environments, for a set of spiking parameters. Experimentally, grid spacings *a*_*g*_ range along the ventro-dorsal axis of MEC from 0.3 m to 1.2 m in smaller environments [3, 12], and from 1.7 m to 3 m and higher in larger environments [14]. The place field sizes grow accordingly, and since the effect of speed is stronger for smaller fields, percolation is least likely to occur in smaller lattices.

For conservative estimates, the grid cell spacing in the simulations was therefore fixed at a lower value, *a*_*g*_ = 60 cm, while the lattice parameter varied from *ξ*_*g*_ = 0.8 (fields almost abut) to *ξ*_*g*_ = 1/3. The spatial phases and orientations of the grid field lattices 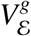 were randomized to represent different possibilities for cells sampled along the ventro-dorsal axis of MEC. To maintain realistic dynamics of spiking activity, the trajectories were generated by re-shaping experimentally recorded paths in open arenas, preserving the observed speed of the animal. The length of each trajectory allowed producing at least 100 path segments extending from one side of the environment to another.

The results show that paths crossing an equilateral triangular enclosure with side *L* = 6 m start percolating grid cells as the lattice parameter and the firing amplitude exceed, respectively, *ξ*_*g*_ ≈ 0.6 and *A*_*g*_ ≳ 20 − 25 Hz, independently from the lattice’s shift and planar orientations (Fig. 2A,B). As *ξ*_*g*_ grows further, percolating paths start appearing at lower firing rates and quickly proliferate, densely covering the navigated area. On the other hand, increasing *A*_*g*_ allows inducing percolation at lower *ξ*_*g*_. The pairs (*ξ*_*g*_, *A*_*g*_) for which the percolation becomes possible form a boundary that separates “percolation phase” from the phase in which percolation is statistically suppressed (Fig. 2B).

**FIG. 2.**
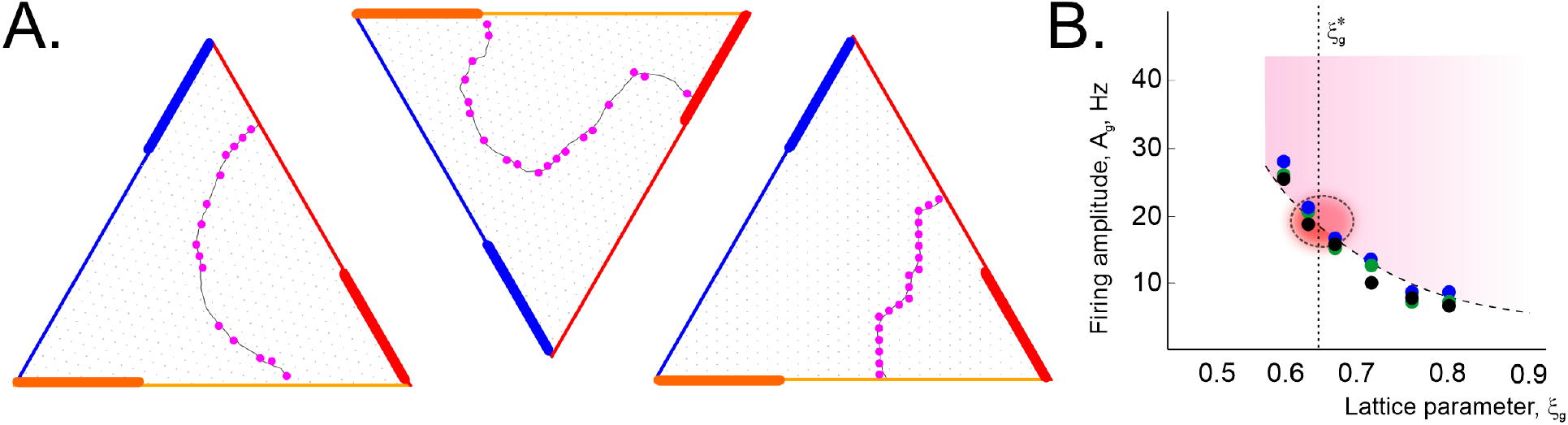
Grid field percolation. **A**. A segment of simulated trajectory (gray line) passing through a large 20 × 20 m triangular environment. Active vertexes are marked by pink circles. The first two paths are non-percolating, the third path percolates the enclosure. Side bars indicate *L*/3 length scale. **B**. For a given set of parameters (*ξ*_*g*_, *A*_*g*_), the onset of the percolation phase was scored when at least 90% of grid cells were percolated by at least 5% of cross-environment path segments, and 90% of paths extending over at least *L*/3 percolate a grid cell. The latter condition defined the size of the simulated grid cell ensemble—about *N*_*g*_ ≳ 200 cells. Dots of different colors mark values obtained using different exploratory trajectories that cover the environment evenly, without artificial favoring one part of the environment over the other. The dashed line separates two phases of the grid cell network’s activity: the pairs of (*ξ*_*g*_, *A*_*g*_)-values on its right (pink area) induce percolation, while the values on its left do not. Encircled area marks the domain of smallest *A*_*g*_ that permit percolation at 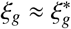.

Furthermore, once emerged, percolation becomes manifested at large scales, e.g., in triangular domains that differ in size by an order of magnitude (Fig. 3A). The effect is strengthened as the cell’s firing amplitude *A*_*g*_ grows, in a manner suggestive of a second-order phase transition controlled by two order parameters, *ξ*_*g*_ and *A*_*g*_ [15, 16].

**FIG. 3.**
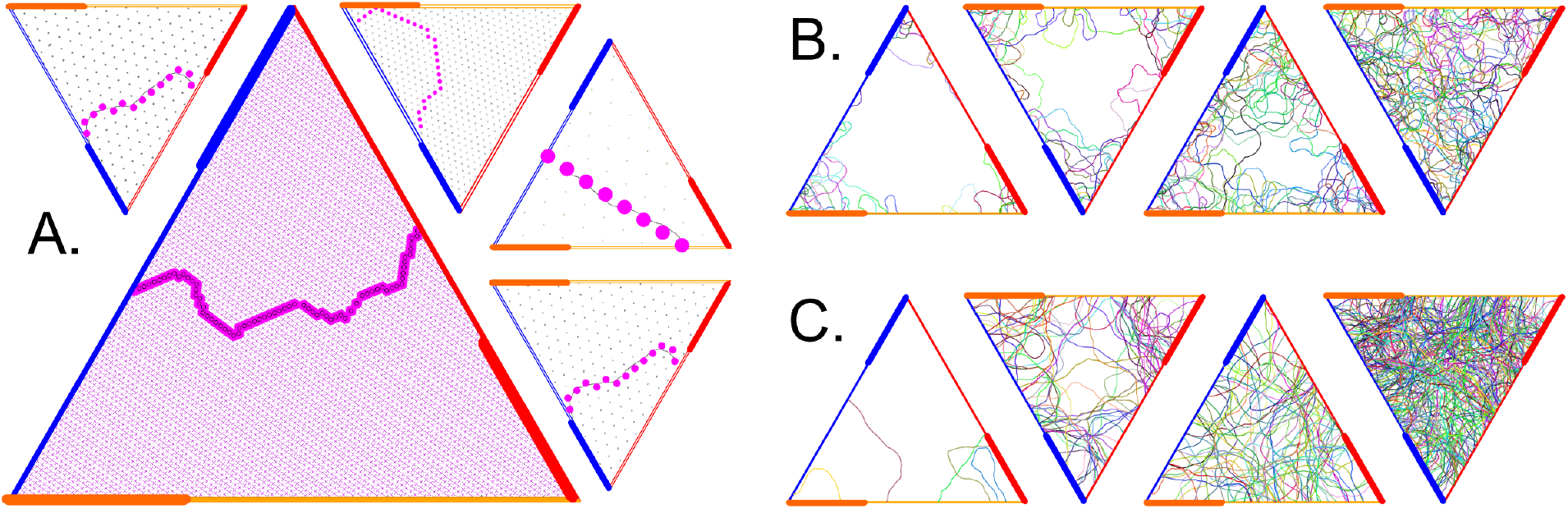
Scaling. **A**. As the lattice parameter exceeds *ξ*_*g*_ ≈ 2/3, percolating paths begin to appear at all scales. Shown are triangular environments with side lengths *L* = 6 m, *L* = 12 m, *L* = 20 m and *L* = 60 m along with a percolating path examples. **B**. As the rate *A*_*g*_ increases from 20 Hz to 200 Hz (*L* = 12 m, *ξ*_*g*_ ≈ 2/3), larger parts of the environment become percolated. **C**. Increasing *ξ*_*g*_ from 2/3 to 0.8 (*L* = 20 m, *A*_*g*_ ≈ 25 Hz) boosts the percolation. Shown are path segments percolating the same cell, each shown with its own color. Side bars indicate *L*/3 length scale.

Note however, that since higher firing rates are energetically costly, electrophysiologically recorded magnitudes *A*_*g*_ should remain close to minimal values that permit a suitable percolation level at a given spatial scale. As shown on Fig. 2B, such estimated values also appear to match the experimental data, which indicates physiological viability of the percolation model.

### Spike lattice in the cognitive map

The results discussed above were obtained by modeling the rat’s moves through the firing fields in the observed environment—an approach that helps visualizing the grid cells’ spiking patterns, but may not directly capture the organization of the underlying network computations [17, 18]. Understanding the latter requires placing the grid cells’ activity into the context of the brain’s own representation of the environment—the cognitive map, encoded, *inter alia*, by the place cell and the head direction cell networks [1, 2]. The computational units enabling this representation are the functionally interconnected groups of hippocampal place cells, *c*_*i*_ [19, 20],

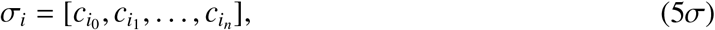

and head direction cells, *h*_*i*_ [21, 22],

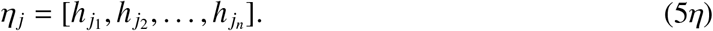

As their constituent place and head direction cells, the assemblies (5) are spatially selective and highlight, respectively, basic locations 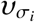 and angular domains 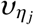 [19–22]. The relative arrangement of these “fields” defines the order in which the assemblies ignite; knowing the latter allows decoding the animal’s positions during active behavior [25–27] and during the “off-line” memory explorations [28–31]. One can hence model hippocampal representation of the grid cells’ firing patterns by the same principle: each individual grid field 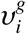 is encoded by those place cell assemblies, 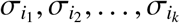, whose fields are contained in 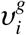, i.e., by the place cells that exhibit coactivity with a given cell *g* and each other.

Computationally, the assemblies (5) are commonly modeled as the cliques of a graph that represents recurrent functional connectivity in the network, e.g., of the *cognitive graph* that represents the collaterals in the CA3 region of the hippocampus [23, 24]. Simulations show that such assemblies form agglomerates, 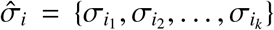, whose joint firing domains, 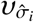, cover the individual grid field regions. The corresponding combinations of place- and grid cells can hence be viewed as the units encoding the *spiking vertexes*,

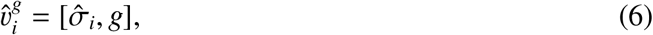

in the hippocampal cognitive map.

The hexagonal order on these vertexes is then established by concomitant activity of head direction assemblies from six “preferred” groups,

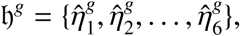

that activate on the runs between pairs of neighboring grid fields, e.g., assemblies from 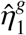 may activate when the rat runs approximately from left to right, assemblies from 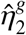 then become active on the runs oriented 60^°^ from the left-right direction and so forth [21]. Correspondingly, the activity of *η*-assemblies from a particular 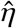-group that leads from a vertex 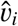 to a neighboring vertex 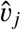 defines an *spiking edge*,

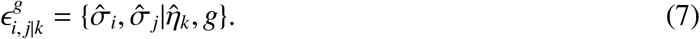

Together, the vertexes (6) and the edges (7) define segments of a *spiking lattice 𝒱*_*g*_ embedded into the cognitive map. In the following, the superscript *g* and subscript *k* will be used to distinguish contributions from different grid and head direction cells, and suppressed otherwise.

### Percolation of spiking lattice

The “intrinsic” definition of the lattice elements (6) and (7) given above leads to a natural generalization of the grid field percolation model. As the animal’s trajectory *γ* traverses a discrete sequence of *σ*-fields,

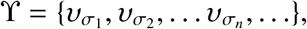

a “firing trace” of ignited place cell assemblies,

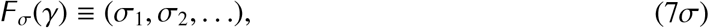

is induced in the hippocampal network, along with a sequence of ignited head direction assemblies,

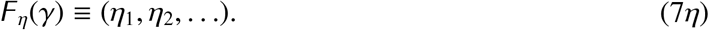

The representation (7*σ*) of the navigated path [25–27] then allows defining *spiking percolation* as follows:

**P1**. A spiking vertex 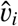 *opens* when its constituent cells activate, i.e., when a place cell assembly from its “hippocampal base” 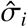 co-activates with the grid cell *g*;

**P2**. Two consecutively opening, neighboring vertexes 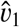 and 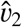 produce an *open spiking edge* in 𝒱_*ℰ*_ if the head direction assemblies from a fixed group 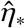remain active on the run from 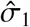to 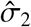;

**P3**. The trace *F*_*σ*_ (*γ*) percolates through 𝒱_*ℰ*_ if it runs through a sequence of consecutively opening vertexes 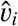 without omissions.

The grid field percolation discussed in §2 can be viewed as a geometric, “pictorial” representation of the spiking percolation if the observed animal moves and the pattern of grid fields are physiologically actualized, i.e., if place cells’ activity marks every location of the rat and if the head direction activity chaperones every move between neighboring grid fields^2^.

Simulations show that the required output is provided by as few as *N*_*c*_ ≳100 active place cells per unit area (1*m* × 1*m*; experimentally observed numbers are higher by an order of magnitude [6]), with typical firing parameters (mean place field size *D*_*c*_ ≈ 24 cm, mean firing rate amplitudes *A*_*c*_ ≈ 20 Hz). Furthermore, *N*_*h*_ ≳ 60 head direction cells firing with the amplitude *A*_*h*_ ≈ 20 Hz over *D*_*h*_ = 20^°^ fields form lattice direction groups 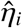 (about 10 cells each) that distinguish runs of the simulated rat between different pairs of neighboring grid fields, which demonstrates that spiking percolation can occur within physiological range of parameters. The resulting series of conjoint open spike edges

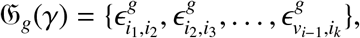

form an intrinsic, spike-lattice representation of the grid field lattice path (1) at the scale defined by the lattice constant *a*_*g*_.

### Path integration

A number of models were built to explain the role of the grid cells in the animal’s capacity to optimize navigation using a cognitive map of ambient environment [34–36]. The mechanisms by which parahippocampal and entorhinal networks learn to represent space and retrieve the obtained information through autonomous network dynamics remain debated [36–38]. A model suggested in [39, 40] implements the required hippocampal replays using persistently firing head direction cells that drive grid cells’ firing from vertex to vertex, which, in turn, activate the corresponding place cell assemblies in spatial order. If the network is trained according to the coactivity between different types of neurons along the navigated path, e.g., 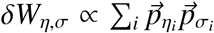, where 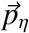 and 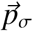 are the population activity vectors, then the learned patterns can be reproduced autonomously in the retrieval phase. For example, head direction firing can be induced by the place cells that start spiking at a position *σ*,

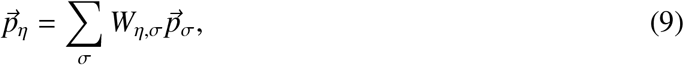

which can then drive the grid cell membrane oscillations, thus generating hippocampal activity at the net step and so forth [39, 40].

From the perspective of this discussion, the network should be trained on the percolating paths only, which can then be reproduced in replays. Aslo note that consecutive activation of two hippocampal assemblies *σ* and *σ*^′^ induced by a persistently firing head direction group *η* may be viewed geometrically as a transition of activity between two adjacent *σ*-locations aligned along the *η*-direction [33]. The Hasselmo model [39, 40] is hence based on using persistent head direction firing to guide place cell activity from a grid vertex to a neighboring one. As it turns out, this mechanism can be generalized to implement the transitions not only along the learned lattice edges, but also to probe their vicinities, which significantly extends the scope of the model.

Consider an edge *ϵ*_*i,i*+1|*k*_ linking two open vertexes, *υ*_*i*_ and *υ*_*i*+1_, along a spike lattice direction 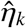. Let 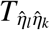 be the matrix permuting the assemblies from two lattice direction groups, 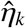 and 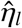 [41]. Then the adjusted weight matrix

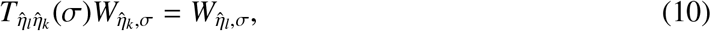

applied at the location *σ* in (9), redirects the persistent head direction activity from 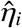 to 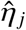 (physiologically, this operation may be interpreted as a cortical or thalamic switch [42]). Two transformations of the weight matrices (10) applied at the ends of an open edge *ϵ*_*i,i*+1|*k*_,

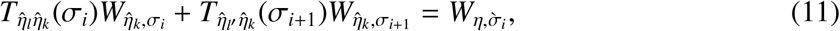

yield the weight matrix that funnels the activity from *υ*_*i*_ and *υ*_*i*+1_ to a side vertex, 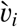 (Fig. 4A). Same mechanism can then reroute the activity from the next open edge, *ϵ*_*i*+1,*i*+2_, to its side vertex 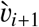, and so forth.

**FIG. 4.**
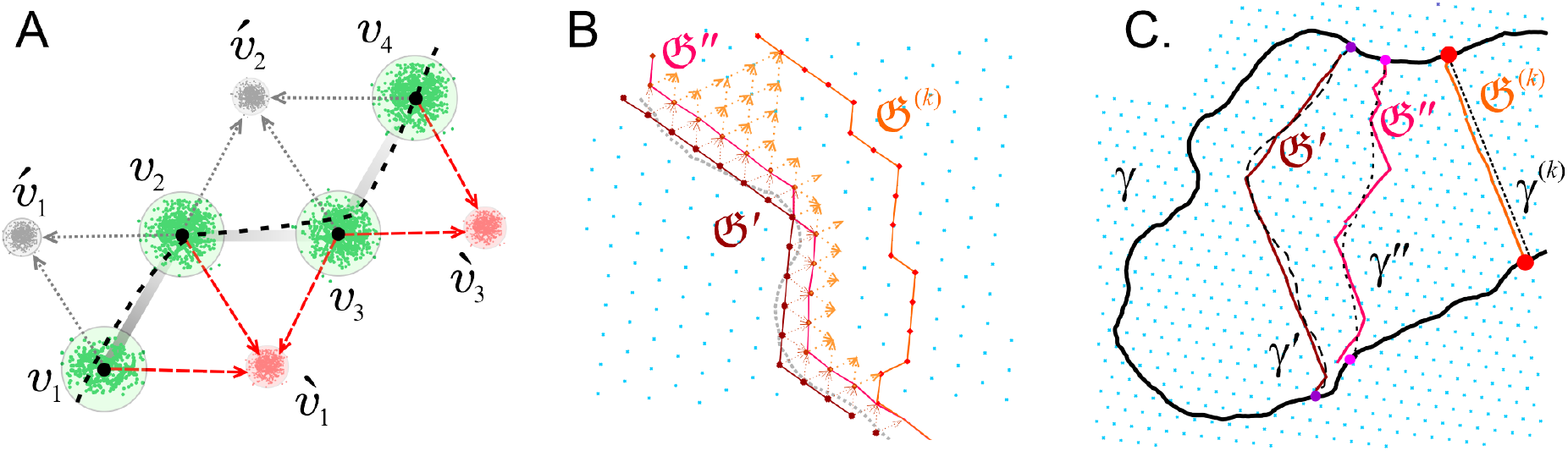
Path deformation. **A**. The activity can transition from an open edge *ϵ*_1,2_ connecting two vertexes *ν*_1_ and *ν*_2_ to its side vertex 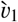 (red field), thus opening the edges 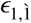 and 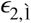 Next, the activity can propagate from the adjacent open edge, *ϵ*_2,3_, to the side vertex 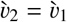, opening the edge 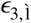, and so forth. The induced shifts are driven towards one side of 𝔊, to allow continuous attractor dynamics. Dashed curve represents a segment of the rat’s trajectory. **B**. A series of transformations (11) can be used to deform the percolating path 𝔊_0_ over the lattice, 𝔊 →→ **C**. Consecutive deformations of discretized paths (𝔊^′^, 𝔊^″^, etc.) can be used to propagate replays of alternative trajectories (*γ*^′^, *γ*^″^, etc.) along the lattice and to produce lattice geodesics, i.e., shortest paths between lattice vertexes, e.g. *γ*^(*k*)^.

From the percolation model’s perspective, activation of the side vertexes also opens the edges that lead to these vertexes, which geometrically amounts to “indenting” the percolated lattice paths (Fig. 4). A series of such indentations can deform and shift the representation of the original path over the spike lattice, 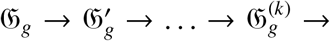…, i.e., induce geometrically deformed lattice paths that can generate hippocampal replays of alternative, “virtual” trajectories and thus guide spatial exploration (Fig. 4B, [43]).

In particular, the possibility of deforming generic percolating paths allows establishing *lattice geodesics*—shortest chains of edges connecting pairs of vertexes. Hippocampal replay of the shortest path between the underlying spatial locations *σ* and *σ*^′^ may account, e.g., for the animal’s ability to run from its current position straight to the nest, which is a key manifestation of path integration [44]. Another implication is that the shortest paths across the spiking lattice define a global spatial metric—the discrete-geodesic distances between pairs of locations [4], (Fig. 4C).

Note that the transformations (11) can be used to redirect the activity to both sides of the open edge series (Fig. 4A). However, if the head direction cells’ firing is to form a single “activity bump” defining a compact range of angles [45, 46], then the activity should be driven to one side of the percolated path 𝔊_*g*_ only. Gradual shifts of the activity bump in the head direction network along a deformed path 𝔊_*g*_ are then consistent with continuous reorientations of the animal’s head.

## III. DISCUSSION

Grid cell activity is commonly studied from the perspective of extracting position codes and spatial metrics from the combinatorics of *ad hoc* defined grid field indexes [5]. Hereby, most models assume, tacitly or explicitly, that a generic grid cell readily conveys spatial regularity of the grid field layouts to downstream networks through spiking outputs, over each navigated path. However, direct simulations show that, over a given traveled route *γ*, most grid cells exhibit irregular spiking patterns that reflect the sequence in which their firing fields were visited, rather than the abstracted order of the fields’ spatial layout. The lattice-like structure of the latter is captured only by those cells, {*g*_1_, *g*_2_, …, *g*_*k*_}_*γ*_ ≡ ℊ_*γ*_, whose grids were percolated by *γ* and which have therefore produced representations, 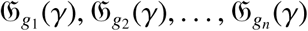, of *γ* in their respective spiking lattices, 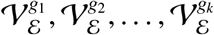. The next path segment, *γ*′, is represented by another percolated group 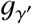 that overlaps with ℊ_*γ*,_ etc. The resulting series of overlapping percolated assemblies form a grid cell firing trace

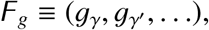

that persistently drive hippocampal activity and allow representing longer, composite paths [*γ* + *γ*^′^ + …]. Note that, from the point of view of grid cells’ operability, the segments *γ, γ*^′^, … may overlap and do not necessarily have to extend across the environment—these assumptions were made above for ease of presentation.

A compact bump of persistent head direction activity can then produce congruous deformations of the percolated path (11) in each contributing lattice, thus generating a compact continuous attractor activity in the hippocampal network [47]. This mechanism allows learning and replaying not only the actual percolating paths, but also their deformations, thus establishing qualitative equivalences between discretized trajectories over spiking lattices, facilitating spatial learning, enabling path integration and defining a global spatial metric of the encoded environment [43].

According to the model, the grid cells’ percolation onset is modulated by the shape of the navigated arena, but it is controlled primarily by several coupled physiological parameters—firing rates, field sizes, lattice spacings, rats’ moves and so forth [15]. Additional restrictions may be required for proper coupling between different cell types, e.g., place field sizes should allow separating grid fields from each other, for encoding distinct vertexes of the spiking lattice 𝒱_*ℰ*_. The full set of conditions defines a *percolation domain* 𝒫 in the parameter space, analogous to the learning region ℒ of parameters required for constructing topologically correct cognitive map from place cell activity [17, 18]. An implication of the model is that the experimentally observed spiking characteristics should fall into 𝒫and allow producing percolating paths in the amounts required for spatial information processing. Certain values can be localized with higher specificity, e.g., the model predicts that the lattice parameter *ξ*_*g*_ should be attuned to the experimentally observed magnitude 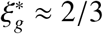, and points at the correct firing rate *A*_*g*_ ≈ 20 − 25 Hz in smaller environments [1, 3–5]. Furthermore, the results point out that changes in one parameter may cause compensatory responses in others, e.g., the network may lower firing rates as *ξ*_*g*_ grows, while producing longer percolating paths at a given lattice scale *a*_*g*_ may require growing *A*_*g*_ or using larger fields, i.e., shifting the grid cell population activity along the ventro-dorsal axis of MEC.

## IV. APPENDIX

**Simulated trajectories** were obtained by reshaping the recorded rat paths and embedding them into simulated environments—triangular enclosures of sizes *L* = 6 m, *L* = 12 m, *L* = 20 m and *L* = 60 m (Fig. 2). The starting position was selected at the boundary of the enclosure randomly, with the velocity directed inward. The trajectory was then generated by time-integrating an experimentally recorded speed series and directing the velocity vector from one wall to another, with random instantaneous deflections distributed over an angular domain [−*α, α*]. The parameter α effectively controls the shape of the trajectory: small αs straighten the paths and larger αs allow more “swirling” curves.

### Site opening probability

The Poisson firing rate of a grid cell *g* is a function of the rat’s position 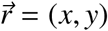

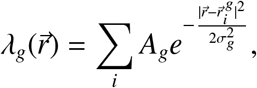

where *A*_*g*_ is the firing amplitude and *σ*_*g*_ defines the size of the firing field 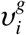 centered at the point 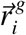. A path segment crossing through 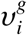 can be approximated by a chord of length *l*, parameterized by the variable *u* and positioned at the distance *l*_⊥_ from the center (Fig. 5A,B). The mean integrated rate of the cell *g* is then

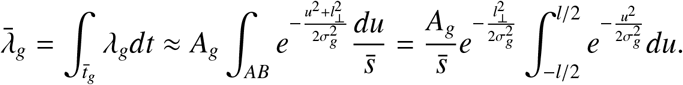

**FIG. 5.**
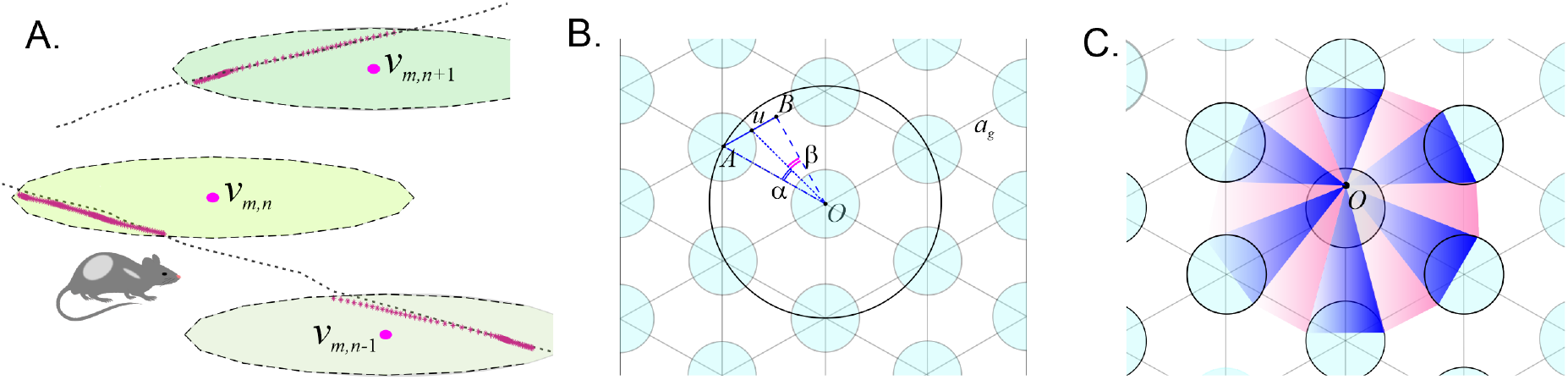
Grid cells. **A**. A segment of the rat’s trajectory can be approximated by a chord cutting through the grid fields. Right panel shows a chord *AB* of length *l* passing at the distance *l*_⊥_ from the firing filed center, *ν*. **B**. The vertex centered at *A* may open if the rat moves within the angular domain *α* (shaded blue); if the rat is directed within the domain *β* (pink shade), then the trajectory escapes. **C**. The geometry of the escape changes as the starting point *O* shifts, leading to small corrections to the probability estimate, proportional to the square of distance between *O* and the field center.

Using *D*_*g*_ ≈ 2 *πσ*_*g*_ for the firing field diameter, and the relationship 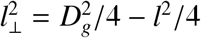 yield

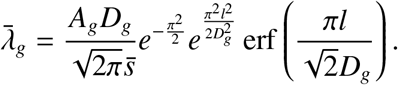

From geometric probability theory, the average chord has length

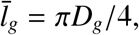

and hence passes at a distance 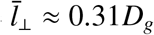 from the field center [11], which allows writing

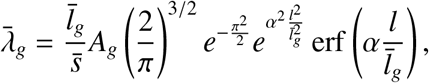

where 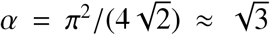. The latter equation implies simply that the mean integrated rate is proportional to the mean time spent to run through the field, 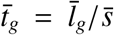. The proportionality coefficient between 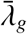 and 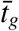 can be interpreted as the characteristic rate during that run,

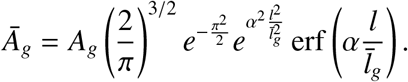

During an average run, i.e., for 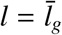,

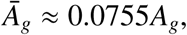

which is equivalent to (4). For example, if the maximal rate is *A*_*g*_ = 25 Hz, then 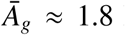 Hz (similar values reported in [14]). If the mean speed is 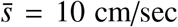 and the mean filed size is *D*_*g*_ = 40 cm, then 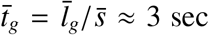 and the net rate is 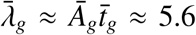, i.e., the cell spikes with probability 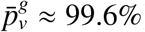). For *A*_*g*_ = 10 Hz, vertexes open with probability 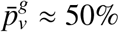.

### Bond percolation probability

Consider the case when the rat moves from the center *ν* of a firing field, outwards along a straight path. The probability *p*_*b*_ of reaching one of the neighboring fields is defined by the ratio of that field’s angular size, as viewed from *ν*, and the angular size of the gap between the firing fields (Fig. 5B). Due to symmetries, it is sufficient to consider the domain bounded by the angle ∠(*AOB*) and the angles *α* ≡ ∠(*AOu*) and 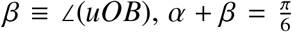, which define the probability as

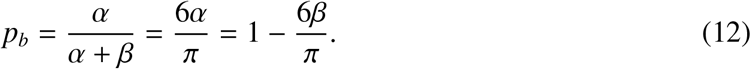

From the lattice’s geometry, |*uB*| = (*a*_*g*_ − 2*R*_*g*_)/2 and 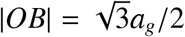. From the triangle *AOu*, the distance 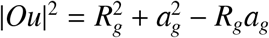, and from the triangle *uOB* one has

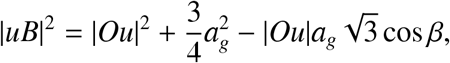

which yields

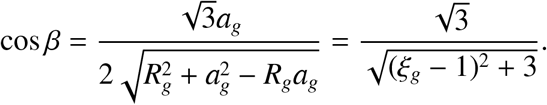

For small lattice parameter, *ξ*_*g*_ → 0 (vanishing grid field size), *β*→ *π*/6, which eliminates the edge opening probability, *p*_*b*_(*π*/6) = 0. Conversely, as the firing field size approaches the gap size, *ξ*_*g*_ → 1, then the gap vanishes, *β* → 0, which leads to the link opening, *p*_*b*_(0) = 1. The physiological value 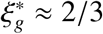 produces *β*^∗^ ≈ 0.19, which corresponds to an overcritical probability, *p*_*b*_(*β*^∗^) 0.637.

If the move starts with an offset Δ*r* from the center of the firing field, *r* = *r*_*O*_ + Δ*r*, then the escape probability (12) will be an analytical function of Δ*r*/*D*_*g*_ ≤ 1. The zeroth-order term in the corresponding (Δ*r*/*D*_*g*_)-expansion is the mean probability given by (12). The first order term will vanish due to symmetries and the non-vanishing corrections are therefore quadratic,

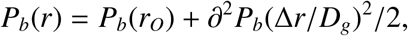

which justifies using (12) for practical estimates.

## Acknowledgments

The work was supported by the NSF grant 1901338.

## Footnotes

1 Throughout the text, terminological definitions are given in *italics*.

2 To simplify modeling, movement direction was used as a proxy for the head direction, although physiologically these parameters not identical [32, 33]

